# Graph Autoencoder and StrNN based Causal Analysis of Mortality in Heart Failure Patients

**DOI:** 10.1101/2024.11.11.622921

**Authors:** Kim Daeyoung

## Abstract

Though analyzed for decades, dissecting and finding mechanisms of cardiovascular diseases, especially heart failures, are still an on-going task for many researchers. However, through recent floods of machine learning and deep learning algorithms to replace traditional approaches, and their applications in diverse cardiovascular research areas, it seems plausible to say that conquering or preventing heart failure catastrophes might no longer be a delusional task within a few more years. To accelerate the arrival of a new era, this research implemented several cutting-edge algorithms currently introduced in causal deep learning to observational heart disease patient data to find key mechanisms that lead to cardiac deaths under a highly flexible framework. Extracting latent causal DAGs from observational data using Graph Auto Encoder, and finding specific causal relationships and interventional effects under Structured Neural Networks (StrNN), novel findings regarding key causes of deaths in heart failure patients were found in numerous aspects. Specifically, existence of intervals where average treatment effects due to causal interventions in platelets, ejection fraction, and serum creatinine levels dramatically decrease or increase was found among heart patients, which can lead to significant eliminations or additions of practical clinical treatments in terms of reducing cardiac death event probability after cardiac failure.

## I. Introduction

**F**or decades, dissecting and conquering the mechanism of cardiovascular diseases(CVD) such as heart failures or myocardial infarctions have been a demanding, yet invaluable task to many researchers. Through numerous efforts in cohort studies and observational studies, key factors that lead to cardiac arrests or mortality due to heart failure, such as serum creatinine levels, serum sodium concentration, glomerular filtration rate or HCM mutations of beta-MHC have been identified and analyzed to prevent death due to cardiac diseases [1-3]. However, most of these works had certain limitations in terms of model usage.

Specifically, majority of research regarding cardiovascular disease commonly implemented cox proportional hazard models [1,4] or logistic regression models [3] to identify the effects of risk factors on CVD, which leads to relatively lower performance in mortality prediction or to false identifications of ground truth effects when highly complex relationships between variables exist in non-constrained environments. Though there were exceptions such as using Poisson regression models [5,6] or temporal reflected logistic regression models [7], conventional statistical models had dominated the overall cardiovascular research for decades, leading to a deadlock in highly advancing prediction performance of mortality or finding complex causal effects that can lead to CVD based deaths in latent aspects.

However, with recent floods of Machine Learning(ML) and Deep Learning(DL) methodologies in data science, it is found in numerous research that this deadlock can be escaped through increased prediction performance acquired from novel frameworks of ML/DL. Moreover, using frameworks from ML or DL, novel indicators or predictors for mortality due to CVD are currently being discovered. For example, body mass index(BMI), which was excluded from important CVD factors in previous research, is being found as a strong critical factor in predicting mortality due to heart failure or in-hospital mortality after percutaneous coronary intervention based on AdaBoost approaches or multi-head self-attention based convolutional neural networks [8,9].

Based on current findings, this research focused on the fact that conquering or preventing CVD based deaths might not be a delusional task within a few more years. To accelerate the era of CVD mortality prevention, this research implemented cutting-edge techniques currently being reported in causal deep learning areas. Implementing novel approaches to find ground truth causal relationships between candidate factors of heart failure under Graph Auto Encoders, and training neural causal models (NCM) using extracted causalities, advanced prediction models for mortality after heart failure and novel findings in interventional effects of influential factors which lead to deaths were analyzed. Results show that under novel deep causal frameworks, prediction performance that surpass ML approaches and intervention effects of causal factors can be efficiently acquired with high validity.

## II. Preliminaries

### A. Related Works

Recently, researches that focus on predicting mortality due to cardiovascular diseases are being conducted under non-conventional methods, such as ML/DL frameworks. Under ML frameworks, decision-tree based methods generally made significant progress in making high performance models for mortality predictions. For example, attribute selection under information gain criterions and AdaBoost usage led to significantly high AUC performance in discrimination between survival/in-hospital mortality, when compared to traditional logistic regression approaches [8]. Here, serum creatinine and BMI were found as novel important features for valid prediction in-hospital mortality after percutaneous coronary intervention (PCI). Furthermore, it was found that implementing Random Forest(RF) methods, boosting classifiers and hierarchical models using RF based prediction probabilities as inputs highly exceed C-statistic performance of traditional Logistic Regression methods regarding analysis of 30 or 180-day readmissions due to heart failure in Tele-HF dataset [10]. Using decision-tree based ML methods for cardiovascular disease research was also found efficient when it was incorporated into neural network based estimations. For example, differentiation performance considering ATH and HCM showed dramatic increase in terms of sensitivity through incorporating speckle-tracking echocardiography to the ensemble of artificial neural networks (ANN), SVM, and random forest [11].

Meanwhile, under DL frameworks, use of deep neural networks itself generally led to higher performance than traditional approaches, even ML approaches. Specifically, in recent deep learning based models, such as FCN, Resnet, or Time LeNet etc., heart abnormality classification accuracy showed striking advancements compared to conventional approaches [12]. Especially under Time LeNet (T-LeNet) usage in processing CHF-RR interval time series data and normal sinus rhythm data, it was found that advanced deep learning architectures can lead to early-stage cardiac arrest or cardiac death preventions. Furthermore, recent multi-head self-attention based convolutional neural networks(DLS-MSM) were shown to make significant improvements in AUCs and F1-scores when predicting mortality in cardiac patients, while SHAP analysis based on DLS-MSM finding pCO2, respiratory failure, BMI, and ARB drugs as key features in predicting mortality within heart failure patients [9].

ML/DL approaches in terms of feature selection or dimensionality reduction in CVD research were also shown effective under diverse performance metrics [13,14]. For example, improvements in classification sensitivity and specificity in traditional models, such as KNN or linear classifiers, were found significant in recognizing congestive heart failure(CHF) based on heart rate variability(HRV) features, when Mutual Information based feature selection methods such as MIFS-U, mRMR, or UCMIFS were used [13-16]. Under these approaches, HRV related NHF(normalized power of high frequency band from RRI) and SDARR(standard deviation of average RR intervals) were relatively found more important than other typical factors. Furthermore, usage of ML models itself such as Support Vector Machines with kernels or eXtreme Gradient Boosting(XGBoost) in feature ranking and feature selections to predict survival of heart failure patients was also found to effectively extract only key indicators such as serum creatinine and ejection fraction for prediction, leading to better results in estimating mortality probability [14].

### B. Neural Causal Models(NCM)

For interventional analysis or effective suggestions regarding practical treatments in medical areas, finding causality between treatments and outcome becomes a prerequisite. Therefore, for deep learning or machine learning to be applicable to practical environments, incorporating causality or injecting causal structures into current ML/DL model architectures is required. One conceptual approach would be using Neural Causal Models(NCM), which is a neural network approach that links Peal, J’s structural causal model (SCM) with deep learning architectures [17,18]. Under theoretical backgrounds and proofs by K, Xia et al. 2022, it was found that when a neural network is constrained by a causal diagram ℊ, by the theorem of NCM ℊ-Consistency and *L*_2_-ℊ representation, deep neural networks under constraint of ℊ can preserve full expressivity and enable canonical analysis in causal inference [17].

One general example, or an initial blueprint of NCM would be a Masked Auto Encoder(MADE) model introduced in 2015 [19]. MADE in causal inference is a ℊ -AutoEncoder model where ℊ is a fully autoregressive directed acyclic graph (DAG), which has a lower-triangular adjacency matrix with all elements under the diagonal being 1. Based on Hadamard multiplication of binary mask matrices with weight matrices, MADE injects ℊ to Autoencoders so that the trained results of MADE can represent structural equations and conditional densities found in ℊ. Though having high representativity, as it only considers fully autoregressive DAGs, MADE has certain limitations in terms of applications for real data analysis environments, where conditional independence due to d-separations are present. To solve this issue, a novel NCM that can incorporate all DAG-type causal SCMs into neural networks was recently introduced in NeurIPS 2023: Structured Neural Networks(StrNN) [20]. StrNN injects domain causal knowledge using binary mask encoders same as in MADE, but enables ℊ to be general DAGs by defining the adjacency matrix as a sparse lower triangular matrix(**Fig 1**). Based on greedy algorithms to find appropriate binary matrix solutions that can maximize the number of links in neural layers under the ℊ -constraint, StrNN was found effective in modelling non-linear, complex causal relations with high performance. Under these characteristics, this research implemented the concepts of NCM, especially StrNN in modelling CVD DAGs to achieve valid causal analysis, and effective mortality prediction for heart failure patients.

**Fig 1.**
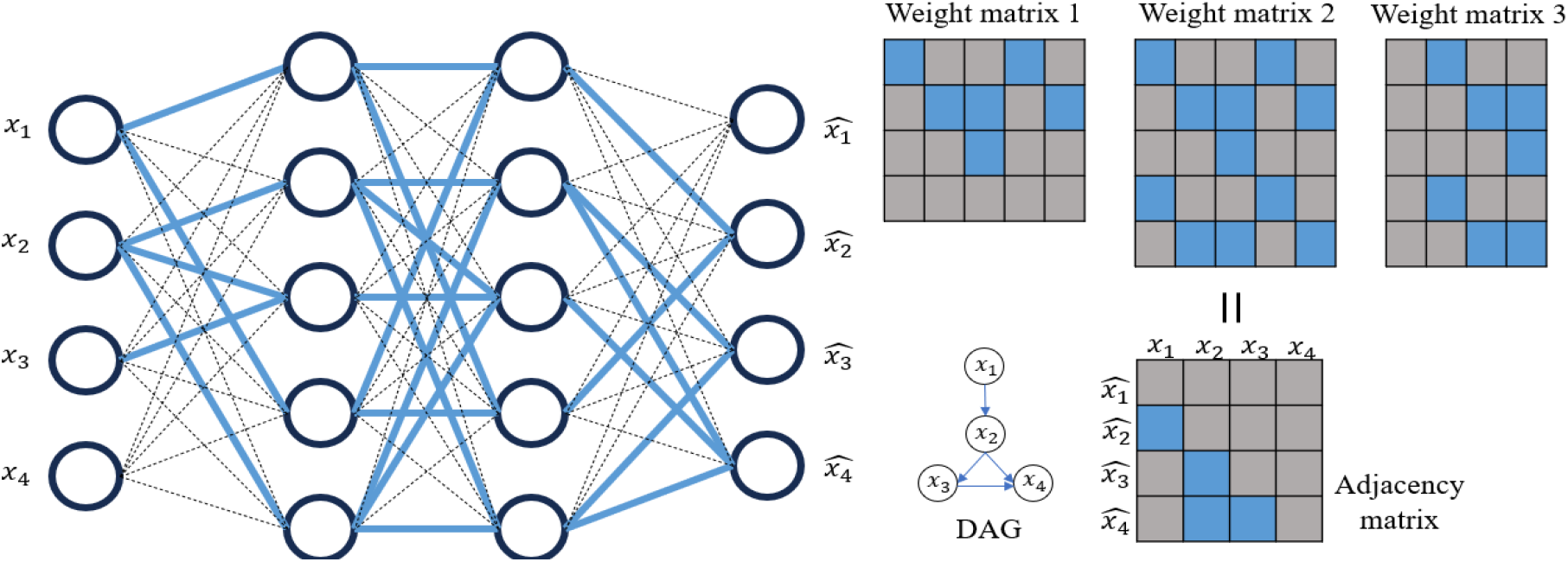
Visualization of StrNN [20] with two hidden layers and causal constraint ℊ(DAG). The result of matrix multiplication should be identical as adjacency matrix. (Chen et al., 2023)

## III. Methodology

### A. Data

Among CVD data regarding mortality, UCI repository Heart Failure Clinical Records data was implemented for research [21]. Based on 299 patients who had left ventricular systolic dysfunction and patients who were classified as class III or IV in New York Heart Association(NYHA) criterion due to heart failure, Heart Failure Clinical Records data contains 13 patient features from studies of Faisalabad Institute of Cardiology and at the Allied Hospital in Faisalabad (Punjab, Pakistan) in 2015 [22]. Specifically, features of patients contain occurrence of death event in follow-up periods of patients, clinical features, such as creatinine phosphokinase(CPK), serum sodium, serum creatinine, platelets, ejection fraction, diabetes, and additional information such as age of patients, smoking status, or length of follow-up periods of patients. Containing significant risk factors found in numerous previous research, Heart Failure Clinical Records data can be used to build effective mortality prediction models, while enabling researchers to find causal relationships between CVD based mortality and risk factors.

Among studies based on this data, one research recently applied ML approaches(RF, GBM, SVM radial) to this dataset and succeeded in constructing high performing death prediction models, while finding the importance of serum creatinine and ejection fraction in predicting mortality of heart failure patients [14]. Based on its findings, this research considered research of [14] as a baseline result when testing the performance of deep causal models for CVD mortality prediction, and checked the existence of relative advantages and novel findings under using deep causal frameworks to predict heart failure patient’s death probabilities. To start with identical settings, ‘time’(follow-up period) variable was excluded from the dataset before modeling [14]. Furthermore, to focus on gender invariant effects, this research also excluded ‘sex’ variable from initial dataset, which led to the usage of 10 explanatory variables and 1 response variable (death event) for model training and testing. After the significance of deep causal modeling being checked, this research then computed causal interventional analysis to find novel, yet adequate treatments or approaches for heart failure patients who are at risk of mortality. (All features were normalized to values between 0∼1 to avoid gradient explosions).

### B. Graph Autoencoder based causal DAG extraction

To find underlying causal relationships among Heart Failure Clinical Records data, extraction of causal DAG between clinical features and mortality was computed via a Graph AutoEncoder framework. Graph AutoEncoders (GAE) [23] are models which find underlying causal relationships between variables by extracting the adjacency matrix of features via encoder-decoder structures under the general assumption of SEM and SCM. Suppose that the ground-truth causal relationship for some feature set × follows the structural equation in (1). As this is a generalized definition of a non-linear DAG, when appropriate functions g1 and g2 are constructed, and convergence in finding parameters for functions: g1 and g2 is achieved, ground truth adjacency matrix *A* can be successfully returned when setting *A*’s components as trainable weight parameters in SEM. In GAE, this process is computed under the process summarized in Fig 2.

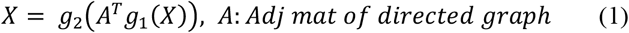

**Fig 2.**
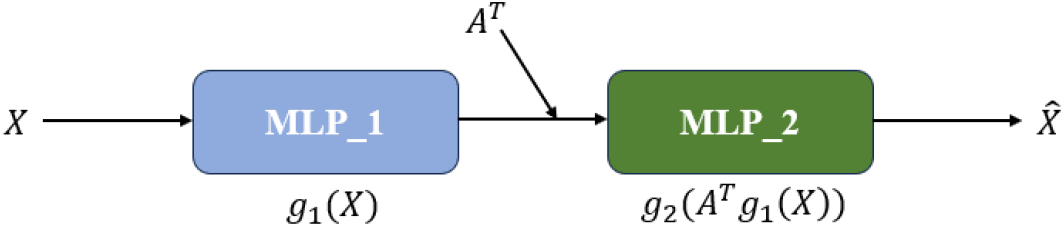
Visualization of Graph AutoEncoder process [23]. First MLP approximates g1, second MLP approximates g2. Weights are updated under the discrepancy between *X* and 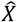.

Using the universal approximation theorem found in deep neural networks, GAE implements multi-layer perceptrons (MLP) to approximate true g1 and g2, which each act as an encoder and a decoder in information transfer aspects. Then, GAE updates parameters in MLP_1, MLP_2 and *A* under the optimization problem formulation of (2) and gradient descent algorithms such as Adam. In optimization problem (2), *θ*_1_ and *θ*_2_ are weight parameters of MLP_1 and MLP_2, λ‖*A*‖_1_ is the regularization term for *A*, 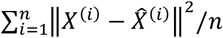 becomes the reconstruction error(MSE) of AutoEncoder outputs, and equality constraint *tr*(*e*^*A*⨀*A*^) = *d* enforces components of adjacency matrix *A* to align with the acyclicity assumption of DAGs. This leads to the use of augmented Lagrange multiplication of (3), with the Lagrange multiplier set as *α*. Based on this approach, parameters *A, θ*_1_ and *θ*_2_ are updated under gradient descent methods, whereas parameters *α* and *ρ* are updated based on iterative formulas summarized in (4). In comparison results with existing methods, such as DAG-GNN and NOTEARs, regarding retrieval of true latent causal links or relationships between variables, this GAE framework was found to be valid and efficient in recovering true DAGs in numerous environments [23-25]. Based on these findings, this research implemented GAE to find causal DAG within features in Heart Failure Clinical Records data.

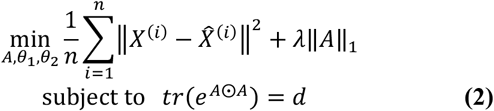

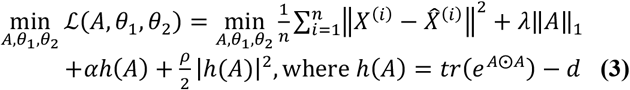

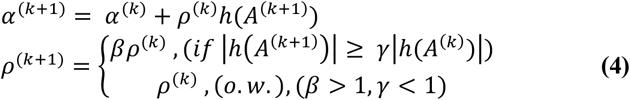

Specifically, this research implemented GAE with an additional assumption for analytic clarity: general additive models (*will denote this approach as GAE-AM: Fig 3). As the equation of (1) does not allow interaction effects between variables, this research attempted to eliminate generation of interaction effects in initial MLP training by setting additive models for each feature in × using weight matrix masking. Similar to ideas of Causal Additive modeling [26], this research allocated 2-layer MLPs to each feature(covariate) to adopt the statistical concept of Generalized Additive Models(GAM), which model dependent structures using the equation defined in (5). In GAM, smooth functions *f*, such as smoothing splines, are used to model diverse dependency structures between response variable and explanatory variables. With neural networks acting as means to approximate an unknown flexible function *f* depicted in (5), in current research, *A*^*T*^*g*_1_ in (1) was approximated with the equation in (6).

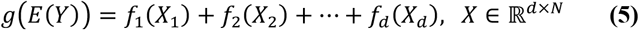

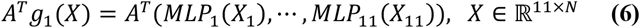

**Fig 3.**
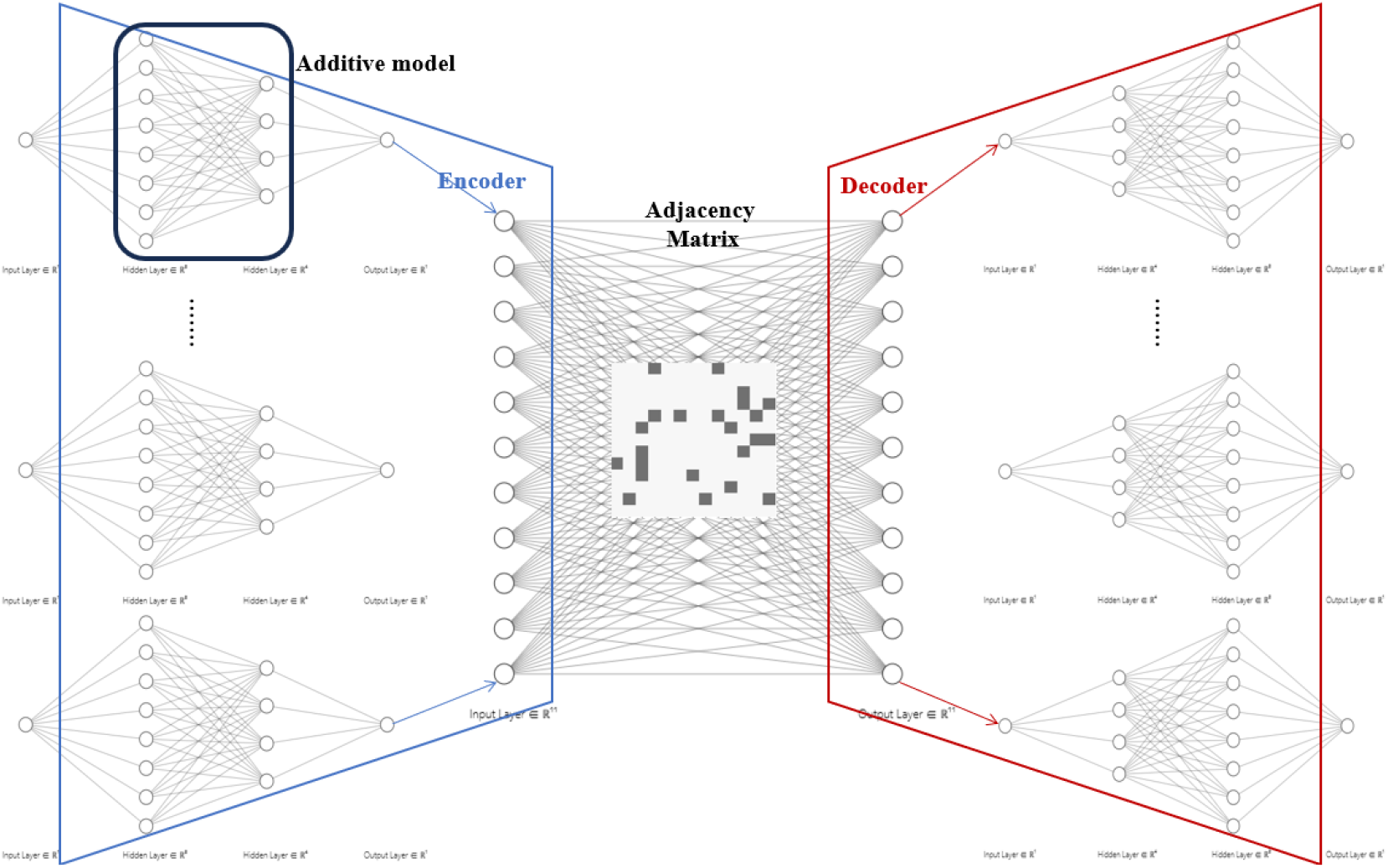
Visualization of GAE-AM. Additive model part depicts 2-layer MLPs for each input variable X. Interaction effects are eliminated through additive modelling.

For advanced GAE usage, this research also implemented a row-wise weighted MSE approach denoted in (7) that can replace Frobenius norm introduced in (2) and (3). Allocating less weight to binary clinical features, while allocating more weight to continuous features, gradient bias generated by AE when using different types of data was efficiently reduced.

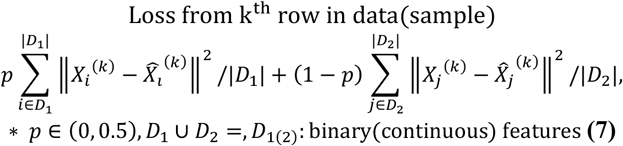

Furthermore, this research added a threshold approach in extracting binary adjacency matrix from GAE-AM. As continuous optimization was implemented for weight updates, the result of trained adjacency matrix becomes a weighted adjacency matrix, which should then be transformed into binary terms so that it can be injected or implemented to diverse Neural Causal Model(NCM) constructions. To ensure high significance in extracted causality, this research assumed a prior hyperparameter *q*, which binarizes adjacency components to 1 if their absolute values are within the 100(1-*q*) percentile from elements in *A* and binarizes components to 0 if values are below the 100(1-*q*) percentile. Therefore, usage of GAE-AM in Heart Failure Clinical Records for causal DAG extraction can be summarized as Fig 4.

**Fig. 4.**
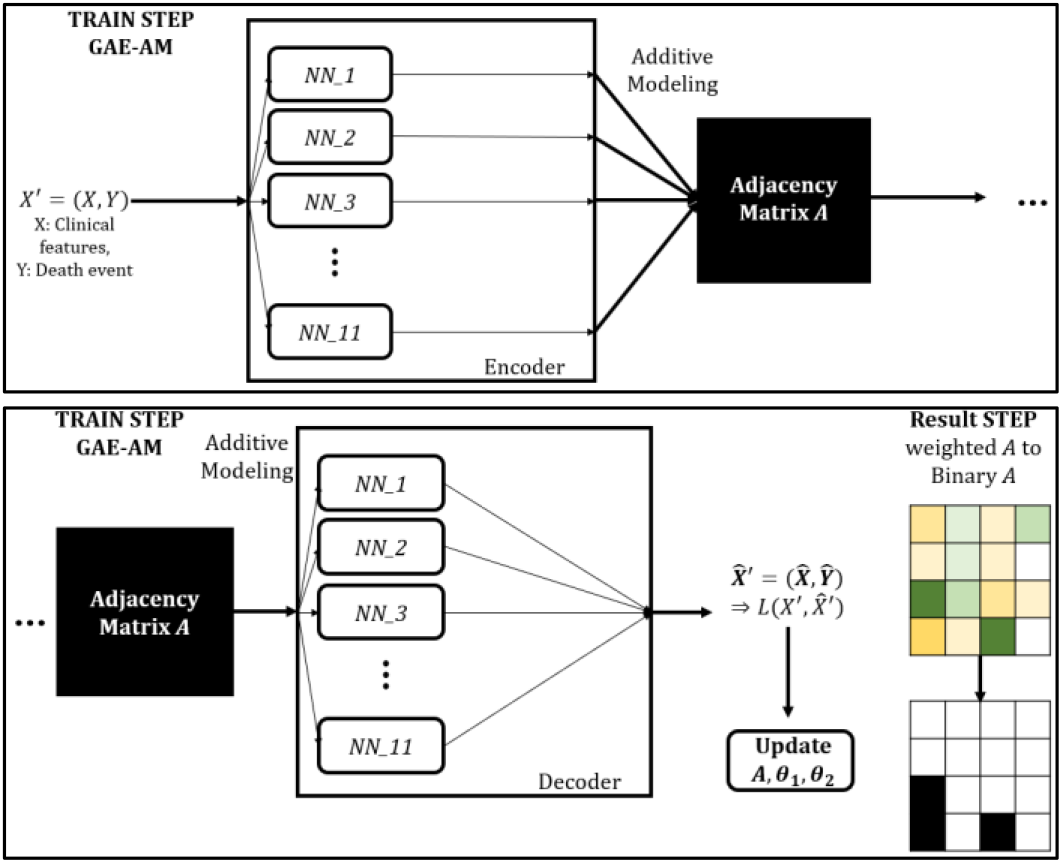
Visualization of GAE-AM process for causal DAG extraction among Heart Failure mortality and general risk factors.

### C. Deep Structured Neural Networks for Mortality Prediction

After implementing GAE-AM to extract causal DAG regarding mortality of heart failure patients, StrNN approach was implemented to check specific forms of causal dependency among components in the extracted network. Setting extracted causal DAG as constraint ℊ in Neural Causal Models, this research constructed a deep Structured Neural Network(StrNN) with four hidden layers and five corresponding binary mask matrices to ensure ℊ-consistency. Here, solving a mask matrix optimization problem denoted in (8) was computed based on the Greedy Factorization algorithm proposed in [20](Table 1).

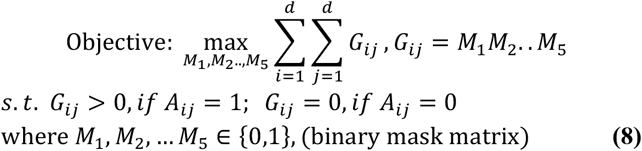

**TABLE I.**
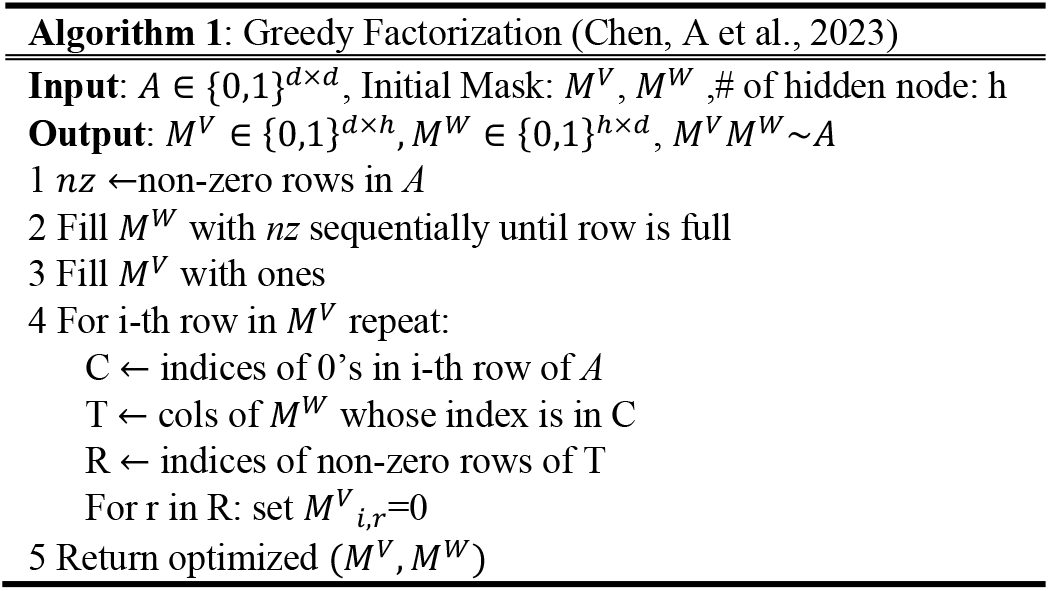
Greedy Factorization algorithm for mask optimization.

To focus more on effects of direct factors on target variable: “DEATH_EVENT”, a 2-hop (from target) subgraph from the extracted causal DAG was set as the adjacency matrix *A* in Algorithm 1. By sequentially repeating the upper process of Greedy Factorization Algorithms for matrices *M*_*i*_(≡ *M*) and *A*_*i*_(≡ *M*^*V*^) for ∀*i*(= 1,2,…,4), which satisfies *A*_*i*_*M*_6−*i*_ ∼*A*_*i*−1_, *A*_4_ = *M*_1_, and *A*_0_ = *A*, individual optimal binary mask matrices: M1 to M5 were returned. These masks were then Hadamard multiplied to corresponding vanilla AutoEncoder weight matrices for ℊ-consistency satisfaction. Through this process, non-linear complex relationships between factors in heart disease and mortality were analyzed, especially causal dependencies between direct causes of mortality (in heart failure patients) and death events.

### D. Intervention Analysis under Do-operators: Interventions in Direct Causes of Heart Patient’s Mortality

Checking validity and effectiveness of StrNN causal models in predicting death events among heart failure patients, when high significance was assured, intervention analysis for direct factors related to mortality was computed to find causal average treatment effects of major risk factors. Here, intervention analysis for each direct factor was driven through do-operators for efficiency. Do-operators (or do-calculus) are methods to check interventional distributions through eliminating all backdoors which are simultaneously linked with treatment variables and an output variable. For example, under Pearl, J’s backdoor criteria and principles of do-calculus[27-29]: (9), when a do-operator is applied to treatment variable × to check average treatment effects on Y under existence of confounder Z, interventional distribution can be deduced as Fig. 5. and (10).

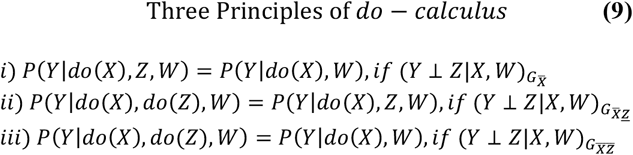

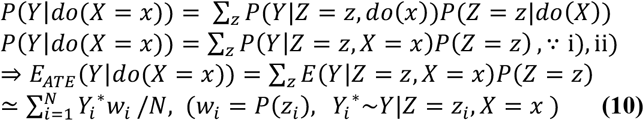

**Fig 5.**
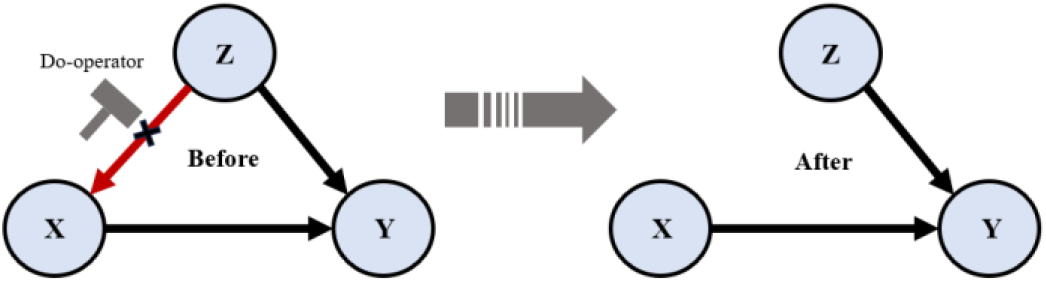
Visualization of Do-operator application in confounder structures. (Red) edge denotes backdoor to treatment variable X.

That is, under the rules of do-calculus and weak law of large numbers(WLLN), the average treatment effect(ATE) of *E*_*ATE*_ (*Y*|*do*(*X* = *x*)) can be approximated by computing the weighted average of samples from *Y*|*Z* = *z*_*i*_, *X* = *x*. Here *z*_*i*_ denotes *i*^*th*^ observation data of Z, and × denotes the intervention on treatment variable X. Based on the conditional distribution of *P(DEATH_EVENT* | *(direct factor))*, which can be extracted from fitting StrNN models, this research sampled 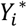 for each possible intervention and multiplied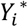 with values of *w*_*i*_, which were computed from density estimations, such as kernel density estimation(KDE).

In this research, estimations of ATE under interventions in direct causal factors to mortality in heart disease patients were checked using the upper approach. After visualizing ATEs of each covariate under samples from StrNN, intervals where drastic ATE increase or decrease takes place were extracted and interpreted to introduce novel treatment strategies that can be easily applied to diverse real-world clinical environments.

## IV. Experimental design

### A. Data preprocessing, and GAE-AM, StrNN settings

Experimental settings for causal analysis(prediction and intervention analysis) in mortality of heart failure patients were set as follows(*Programming language set as Python). For data preparation, the total Heart Failure Clinical Records dataset was randomly divided into train data and test data with proportion of 6(train):4(test) under train_test_split() function in sklearn.model_selection. Proportion of 6:4 was selected to check robustness of causal inference, when compared to the baseline research of [14], which used 7:3. After splitting data, for each sub-dataset, data preprocessing based on normalization was computed to avoid gradient explosions in further analysis. For GAE-AM and StrNN constructions, Tensorflow libraries and Tensorflow Keras functions were implemented for high reproducibility and efficiency (Tensorflow version: v.2.12.0). For applications of mask matrices in GAE-AM and StrNN, custom constraints, which return Hadamard multiplication values of weights, were generated by creating a class that inherits tf.keras.constraints. Constraint [30] and then were applied to kernel_constraint (attribute) in keras.layers Dense().

In GAE-AM, MLPs with 2 hidden layers(8 nodes for first hidden layer, 4 layers for second hidden layer) were implemented as individual additive models for each input variable, thereby leading to an encoder and a decoder structure which has 88 nodes for first hidden layer, and 44 nodes for the second hidden layer. For activation functions in GAE-AM, ‘linear’ function was implemented except for the output layer in the decoder, which was set as having a sigmoid activation function due to normalization of data. For weight initialization methods, He-normal initialization [31] was implemented for high efficiency in gradient updates. Meanwhile, to construct the adjacency matrix multiplication part in GAE-AM, an additional FC-layer with dimension ℝ^11×11^ was attached to the front of the decoder to represent individual components of the adjacency matrix and to update each component under continuous optimization. For weighted MSE loss function settings, parameter *p* was set as 0.3 to avoid high gradient imbalance between binary variables and continuous variables.

After construction of GAE-AM structures being complete, specific parameters to train GAE-AM were selected as in Table II under light manual tuning. Here, masking the last row within the adjacency matrix layer weight matrix, which works as a means to black list all edges starting from DEATH_EVENT (*impossible under time sequence consideration), was implemented. Therefore, the optimization objective introduced in (3) was altered into a new objective in (11).

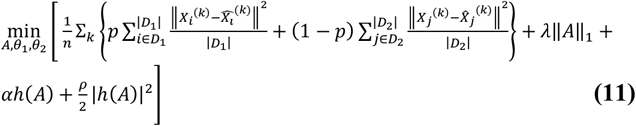

**TABLE 2.**
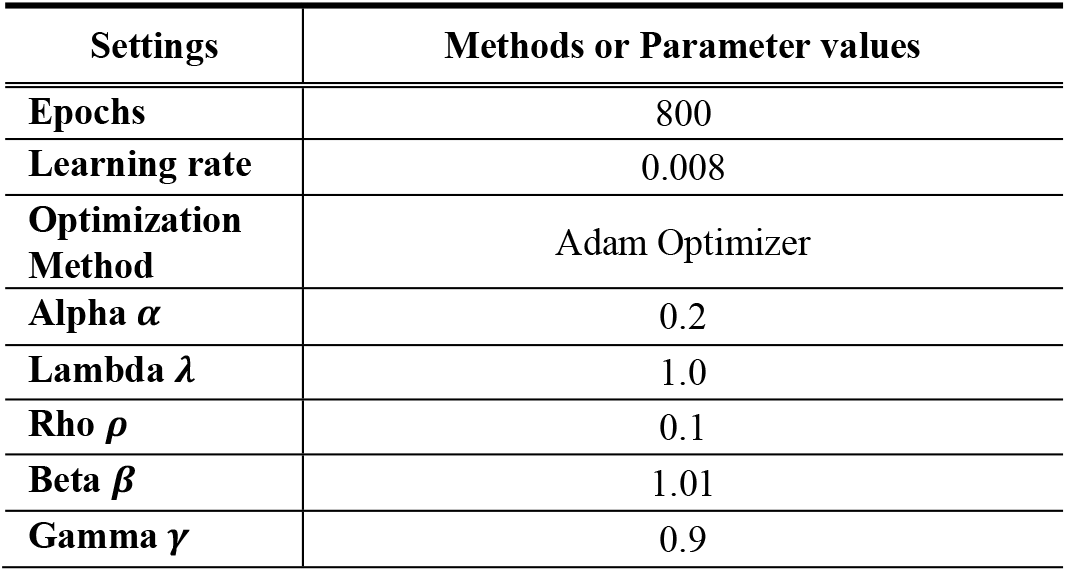
Settings of training methods and parameters in GAE-AM.

After fitting(training) was completed using train data, resulting weighted-adjacency matrix *A* was extracted from GAE-AM. For binary threshold parameter *q, q*= 0.15 was selected to extract only highly significant causal relationships between variables. Under binarized adjacency matrix, directed acyclic causal graphs were visualized and analyzed with the *networkx* python package.

For StrNN implementations, 2-hop subgraph from extracted causal DAG was injected to the following vanilla AutoEncoder structure: 4 hidden layers with {30, 20, 20, 30} nodes, Glorot(Xavier)-Uniform weight initialization [31,32], sigmoid activation functions for each layer and a MSE loss function with reconstruction error from nodes with no parent nodes masked as 0. Furthermore, optimal binary mask matrices deduced from greedy factorization in TABLE I were applied to the AutoEncoder structure through the same procedure explained in GAE-AM. Due to prohibitions of self-loops in DAG, DEATH_EVENT data was not passed into hidden layers from input, and for parent nodes which only have child nodes, hidden layer information was not passed to corresponding output nodes. In StrNN training, same train data, which was used in the process of training GAE-AM, was re-implemented. For optimization methods, Adam optimizer with learning rate 0.01 was used for efficiency, with number of epochs set as 2000 and mini-batch size set as 20 and 50 (two different models checked). With convergence being checked, after StrNN training, fitted StrNN’s prediction performance was checked using test data.

Specifically, prediction performance regarding morality in heart disease patients was tested under four metrics: accuracy, F1-score, ROC AUC score and Matthews Correlation Coefficients(MCC). Furthermore, to check relative advantages of implementing deep causal frameworks in CVD research, comparison with previous baseline performances reported in ML-based research [14] (under same data) was driven.

### B. Intervention Analysis settings

After validity being checked in fitted StrNN, this research implemented intervention analysis regarding factors that directly affect death events in heart failure patients. For interventions in treatment variable X, 100 sequential values from python’s numpy.linspace(0,1,100) were used. That is, 100 sequential values in ascending order between 0 and 1 were used as hypothetical inputs to StrNN. For other influential variables: *Z*_*i*_, such as confounders, real observation values were used as input to implement do-operations. Lastly, for *w*_*i*_ which was introduced in (10), densities of variables were estimated through KernelDensity(bandwidth=0.5) functions in python sklearn.neighbors package.

To check estimated ATEs for each treatment intervention, prediction values for variable: “DEATH_EVENT” from fitted StrNN were used. That is, from the resulting vector ŷ, which equals *StrNN*(*do*(*X* = *x*), *z*_*i*_)), ŷ_*DEAT EVENT*_ was set as 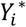 in (10). Through computation of ATEs under approximation in (10) and visualization of ATEs through line plots, novel clinical characteristics, such as existence of intervals where drastic ATE increase/decrease occurs or existence of intervals where ATEs are invariant to interventions, was checked and analyzed.

## V. Experimental results

### A. GAE-AM fitting results

Results for GAE-AM fitting were computed as follows. Fig. 6 and Fig. 7 each denote loss function values per epochs and the returned weighted adjacency matrix under GAE-AM fitting. In figures of loss function in Fig 6, successful convergence to 0 in loss values, which are based on GAE-AM weights(*A, θ*_1_, *θ*_2_), was checked. (Final Weighted MSE = 0.0879, Causal Loss = 0.1425). This implies that the resulting adjacency matrix from 800 epochs can be considered as a valid representation of the underlying causal SCM within Heart Failure Clinical Records data. Furthermore, as visualized in Fig. 7, successful incorporation of non-acyclicity conditions in optimization was found in final steps (epoch=800, non-diagonal elements reduced to values close to 0, Index 0 to 10 (sequentially) denote age, anemia, creatinine phosphokinase, diabetes, ejection fraction, high blood pressure, platelets, serum sodium, smoking and death event).

**Fig 6.**
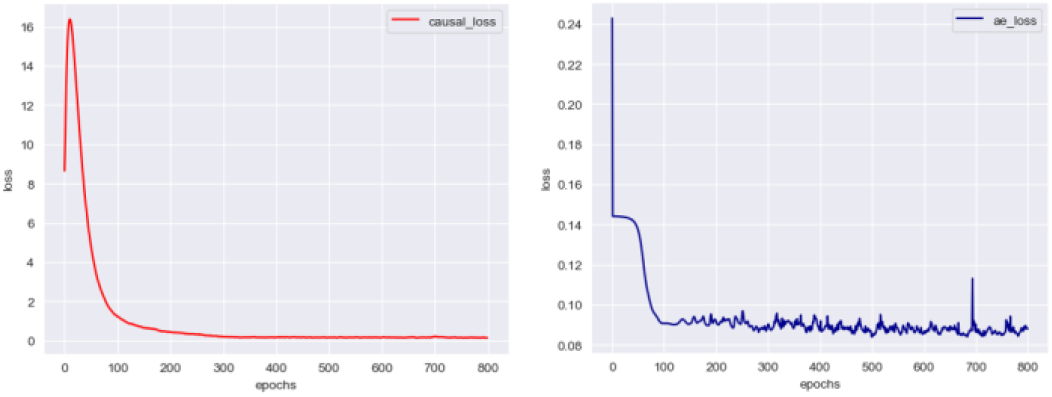
Visualization of Loss function values in GAE-AM. (Left) Causal Loss denoted in (11), (Right) Sole Autoencoder Loss: only Weighted MSE part in (11)

**Fig 7.**
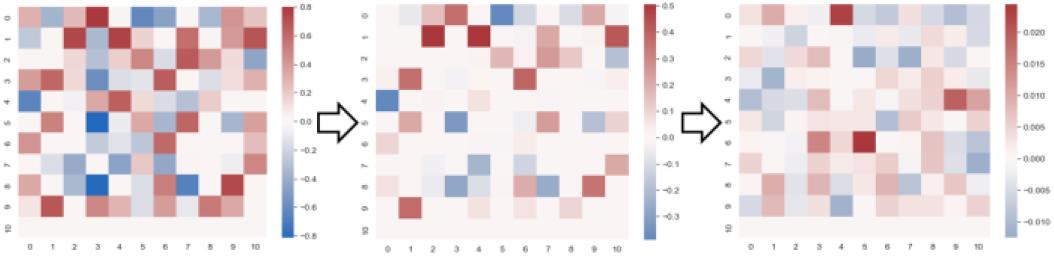
Visualization of weighted adjacency matrix updates in GAE-AM fitting. From left: 10 epochs, 100 epochs, 800 epochs.

Based on high convergence in fitted GAE-AM, binary adjacency matrix (based on parameter *q*=0.15) was computed from the returned weighted adjacency matrix *A*. Results were deduced as Fig. 8.

**Fig 8.**
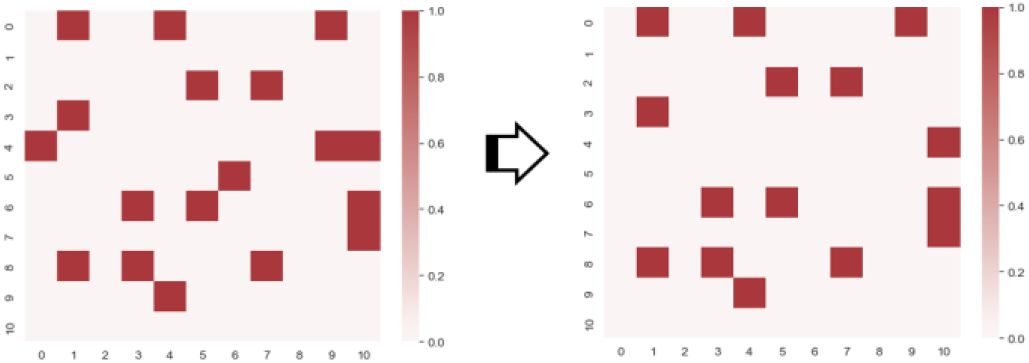
Visualization of binarized adjacency matrix. (Left) Initial binary adjacency matrix. (Right) Modified binary adjacency matrix

Among converted elements in binary matrix, though the vast majority of elements were found not to violate DAG assumptions supposed in experimental settings, three cases were found to violate the directionality assumption in DAGs. Based on the inference that this is a result of minor fitting errors due to weak constraints from implementing the Augmented Lagrange multiplication approach, discarding one of the bidirectional edges within three cases under temporal precedence consideration and domain knowledge was implemented (\**A*_4,_, *A*,, *A*, set as 0). This research therefore implemented the final right adjacency matrix in Fig. 8, which incorporates all error modifications, for further causal analysis.

### B. StrNN fitting results

Results of fitting StrNN models based on the binary adjacency matrix deduced in GAE-AM were as follows. First, extraction of 2-hop subgraph from extracted causal DAG was implemented (Fig 9). After subgraph extraction, this subgraph was injected as constraint ℊ in StrNN construction. Through fitting, training loss and metric(MAE) values for each epoch were deduced as in Fig 10, which both showed successful convergence as the number of epochs increased. (Final MSE=0.5183, MAE=1.3435 under batch size=50). This implies that StrNN successfully extracted non-linear relations between causal risk factors and mortality in heart failure patients. Based on fitting results, this research then checked death event prediction performance of StrNN using test data. Performance metric values based on train/test data were denoted in TABLE III. Though target data was imbalanced(survival: 203, death: 96), StrNN showed significantly high accuracy and f1-scores in predicting mortality among patients who had left ventricular systolic dysfunction and patients who were classified as class III or IV in NYHA criterion due to heart failure.

**TABLE 3.**
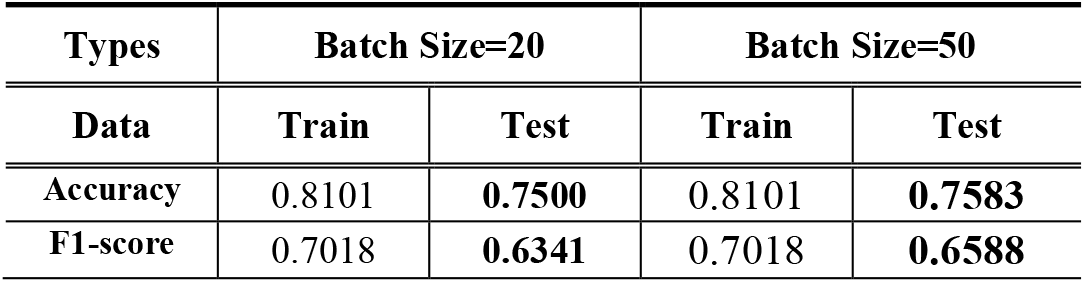
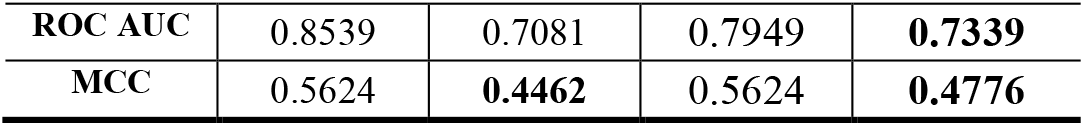
Performance metrics for StrNN(train, test)

| Types | Batch Size=20 |  | Batch Size=50 |  |
| --- | --- | --- | --- | --- |
|  | Train | Test | Train | Test |
| Accuracy | 0.8101 | **0.7500** | 0.8101 | **0.7583** |
| F1-score | 0.7018 | **0.6341** | 0.7018 | **0.6588** |
| ROC AUC | 0.8539 | 0.7081 | 0.7949 | **0.7339** |
| MCC | 0.5624 | **0.4462** | 0.5624 | **0.4776** |

**Fig. 9.**
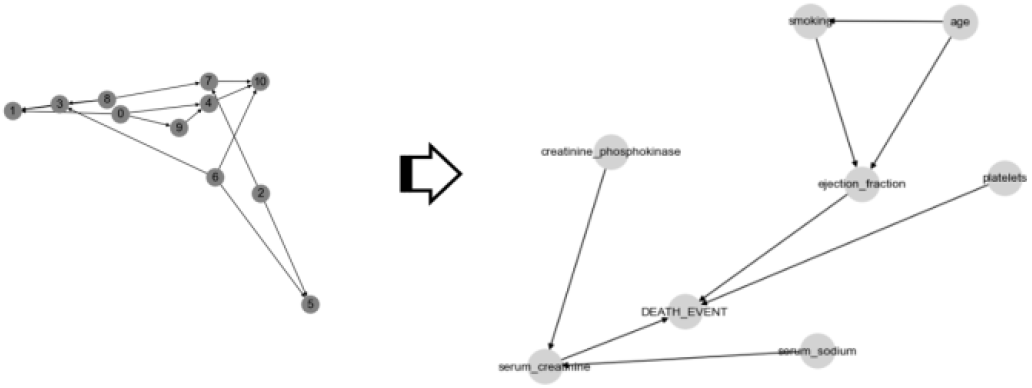
(Right) 2-Hop subgraph(sub-DAG) from extracted causal DAG. (Left) Causal DAG extracted through GAE-AM. Platelets, ejection fraction and serum creatinine were found as direct causal factors for death events in heart failure patients.

**Fig. 10.**
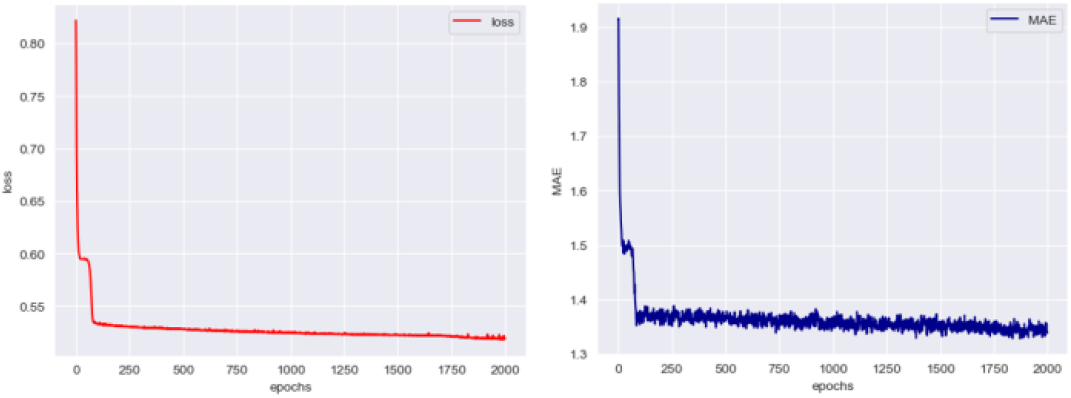
Visualization of Loss and Metric values in StrNN(batch size=50). (Left) weighted MSE Loss, (Right) weighted MAE metric.

**Fig. 11.**
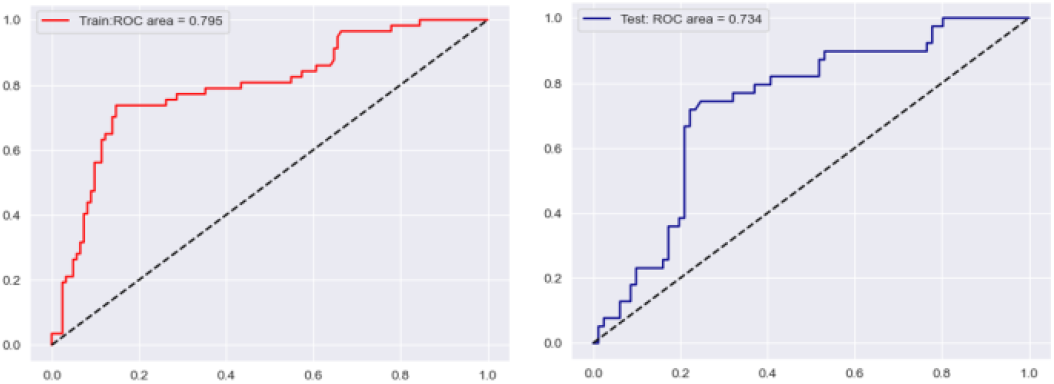
StrNN(batch size=50) based ROC AUC curves for train data(left) and test data(right). High performance in mortality prediction checked.

Moreover, performance of StrNN based predictions highly out-performed performances found in ML based baseline research[14](*introduced *III. Methodology. Data*). For example, in baseline research, when ML based feature selection was implemented and was used to train prediction models based on RF, GBM and SVM radial methods, highest accuracy value and MCC among models were computed as 0.585 and 0.418. That is, StrNN with batch size=50 outperformed baseline results in test accuracy and MCC by approximately 17.3%p and 5.96%p. Furthermore, StrNN results also exceeded general prediction performances resulting from entire feature set usage in baseline research. Regarding entire feature set usage in baseline research, best performance values for accuracy, F1-score, MCC and ROC AUC were sequentially deduced as follows: 0.740 (Random Forest based), 0.554 (Decision Tree based), 0.384 (Random Forest based) and 0.800 (Random Forest based). This shows that though significantly less data was used, test accuracy, f1-score and MCC from neural causal modeling (StrNN) still highly outperformed results from ML based modeling. Thus, results imply that through StrNN, more accurate and reliable predictions regarding death in heart failure patients can be derived, when compared to ML based approaches. Moreover, unlike ML model based research, which can only check correlations between variables, using NCMs such as StrNN can lead to analysis of causal relationships between variables. This implies that strong advantages of using NCM frameworks in CVD prediction or risk factor analysis exist, compared to traditional CVD analysis methods.

### C. Intervention Analysis

Based on high performance found in fitted StrNN(batch size=50), this research finally computed intervention analysis based on direct causal factors of death events from the extracted causal DAG in Fig. 9: Serum Creatinine(src), Platelets(plt) and Ejection Fraction(ef). Based on do-calculus rules dealt in (9) and (10), interventional distributions to calculate ATE for each factor were computed as (12) ∼ (14) (DE: DEATH_EVENT). After acquiring interventional distributions for each factor, ATE was estimated under StrNN(batch size=50) based *Y*^*^sampling.

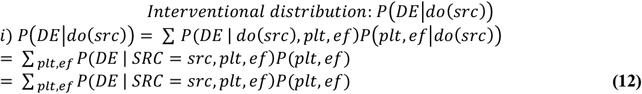

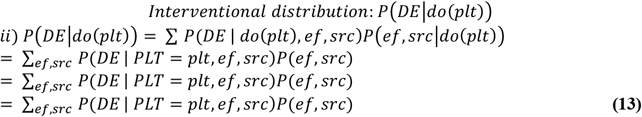

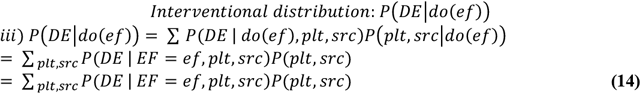

Resulted (estimated) ATEs of direct factors: serum creatinine, platelets and ejection fraction, were visualized in Fig 12-14. In serum creatinine intervention analysis, positive(+) causal intervention effect was found to exist between risk of mortality and levels of serum creatinine in interval [0mg/dL, 4mg/dL]. On the other hand, in interval [4mg/dL, 8mg/dL], invariance to interventions was found in ATEs. This implies that treatments to lower serum creatinine levels in patients who have serum creatinine levels that highly exceed 4mg/dL have incremental or almost no advantage in reducing the possibility of death events in heart failure patients. However, for patients who have serum creatinine levels that are below 4, especially in interval [1mg/dL,2mg/dL], treatments to lower serum creatinine levels can lead to drastic reductions in possibility of mortality. This implies that in serum creatinine clinical treatments, (interval based) cluster-wise approaches should be driven separately to increase the likelihood of survival in patients who suffered from heart failures. This is a significant finding, as previous works only emphasized the existence of positive causal relationships between serum creatinine levels and CVD severance itself [1,33,34]. By finding the existence of a change point in ATEs, inappropriate clinical approaches, such as means to reduce serum creatinine levels within the range of [4mg/dL, 8mg/dL] in heart failure patients, can be avoided.

**Fig. 12.**
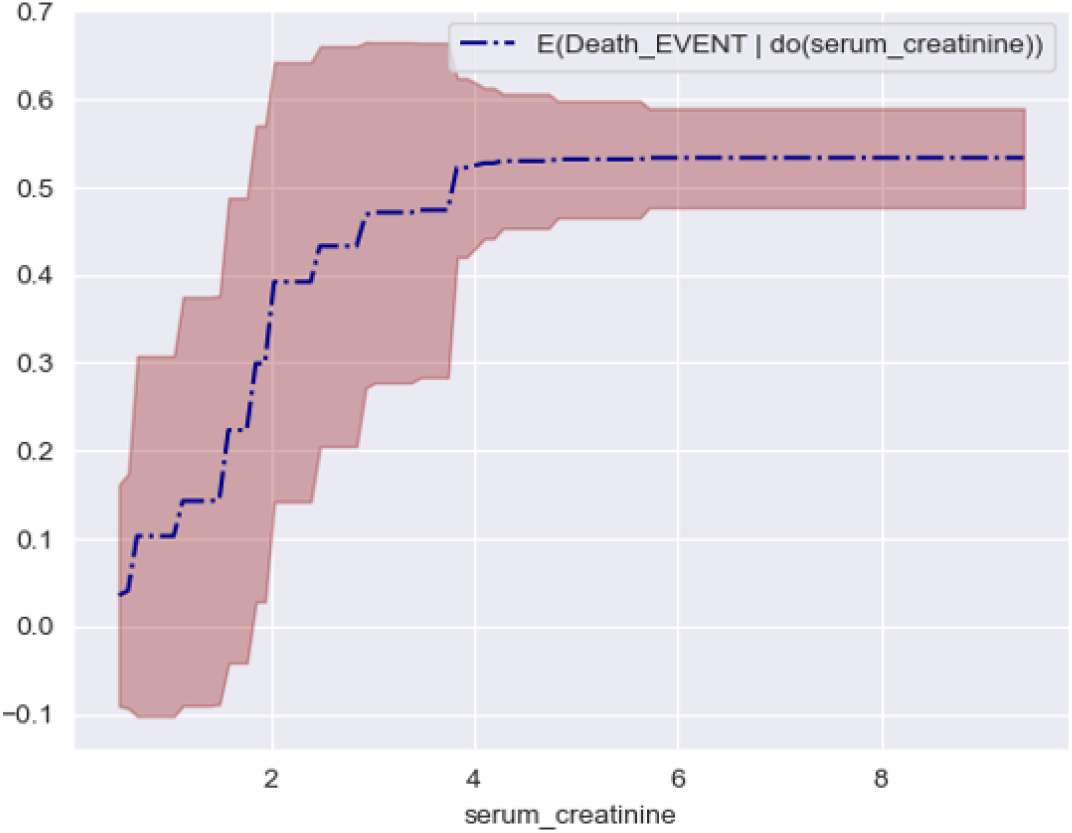
Visualization of Estimated ATEs of serum creatinine on death event in heart failure patients. Colored area denotes of *Y*_*i*_^*^*w*_*i*_ values. Dotted line denotes estimated ATE.

**Fig. 13.**
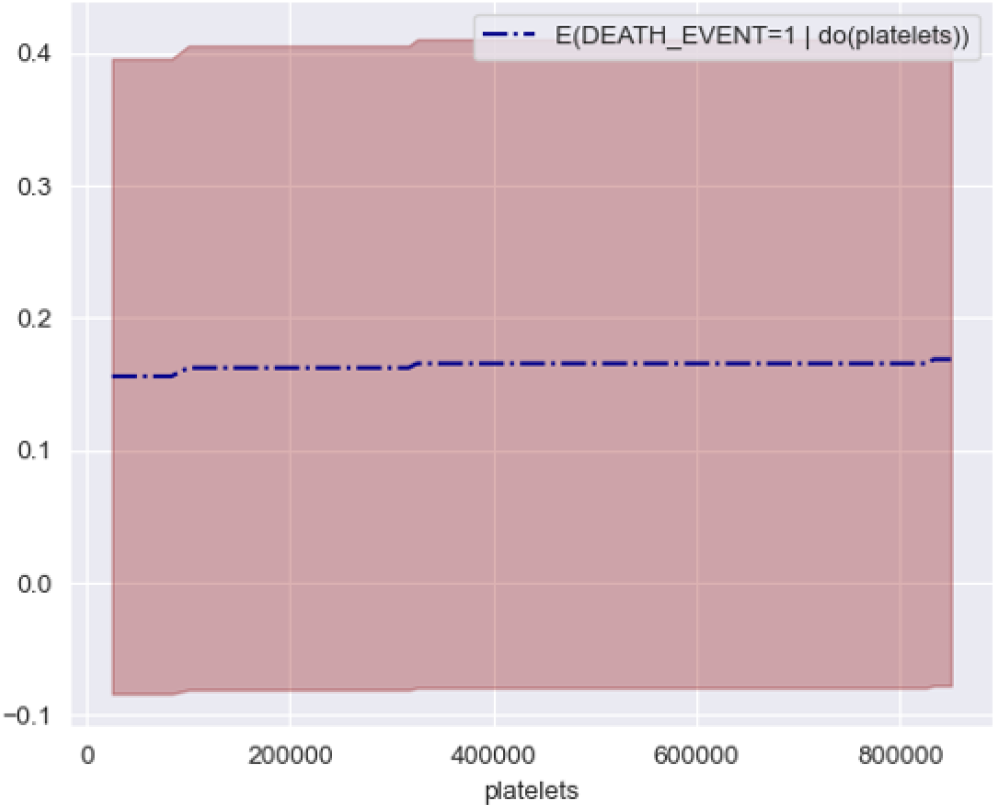
Visualization of Estimated ATEs of platelets on Death event in heart failure patients.

**Fig. 14.**
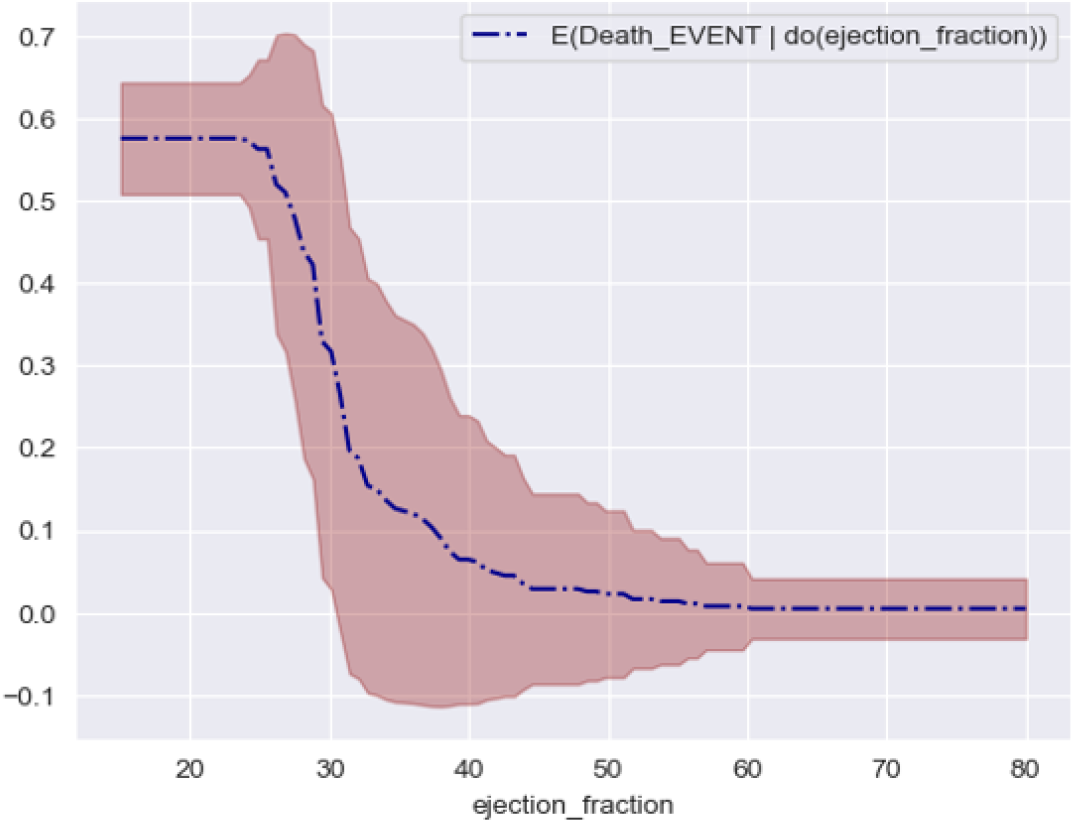
Visualization of Estimated ATEs of ejection fraction on death event in heart failure patients.

Meanwhile, in platelet intervention analysis, positive(+) causal intervention effects were found to exist in multiple intervals. That is, monotonic increase in ATE was found across all intervals in platelet interventions. For example, ATE increase regarding death probabilities of heart failure patients was found to be invariant to interventions within intervals: [0 kiloplatelets/mL, 80000 kiloplatelets/mL], [100000 kiloplatelets/mL, 320000 kiloplatelets/mL] and [340000 kiloplatelets/mL, 800000 kiloplatelets/mL] (*approximate), whereas interventions between these three intervals were found to increase the risk of mortality. This implies that though an excessive amount of platelets does lead to an increase of mortality, in the aspect of treatments, it is hard to expect drastic improvements in death probabilities due to the existence of multiple invariant intervals, which leads to a monotonic decrease in risk of mortality among heart failure patients.

Lastly, in ejection fraction intervention analysis (Fig 14), three distinct ATE intervals were found to exist: [0%, 25%], [25%, 45%] and [45%, 80%]. Though general findings, which states that ejection fraction negatively(-) affects CVD, were also found valid in this research, specific ATE characteristics were found to be different based on types of intervals. In interval [0%, 25%], ATE was found to be invariant to interventions in ejection fraction. That is, heart failure patients who were reported to have ejection fractions below 25% may still have a high risk of death even though certain treatments to enhance ejection fractions were implemented. On the other hand, in interval [25%, 45%], interventions to increase ejection fraction were found to significantly (drastically) decrease the probability of mortality in heart failure patients. This implies that the survival probability of patients who have low ejection fraction levels between 25% and 45% may have significant advancements in terms of remissions or recovery in heart failure when treatments to enhance ejection fractions are applied. In interval [45%,80%], however, invariance to interventions were found as in interval [0%, 25%]. Therefore, clinical treatments which try to increase patient’s ejection fraction within interval [45%, 80%] may have incremental or no effect in decreasing death risk probabilities among heart failure patients.

### D. Intervention Analysis regarding indirect factors

To further expand NCM based heart failure factor analysis beyond direct factors of Death Events, this research also computed intervention analysis regarding two indirect factors of death events in heart failure patients: smoking and creatine phosphokinase. Based on do-calculus rules dealt in (9) and (10), interventional distributions to calculate ATE for each factor were computed as (15) and (16). *In (15), backdoor path to ejection fraction created by factor: ‘age’ was eliminated through do-calculus.

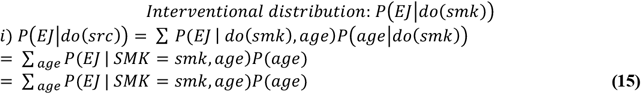

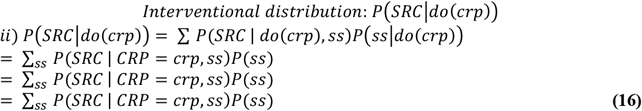

Results of ATE estimations regarding indirect factors: smoking and creatine phosphokinase were deduced as Fig 15. and Fig 16. In smoking interventions on ejection fraction of heart failure patients, a negative(-) causal relation was found between two factors. As smoking was intervened from 0 to 1, estimated ATE on ejection fraction exhibited decrease from 32 to 30. This implies that though being an indirect causal factor of mortality in heart failure patients, intervening on smoking can have significant effects in alleviating the possibility of death events, when considering the intervention effects of ejection fraction found within the interval: [25%, 45%] in *C*.

**Fig. 15.**
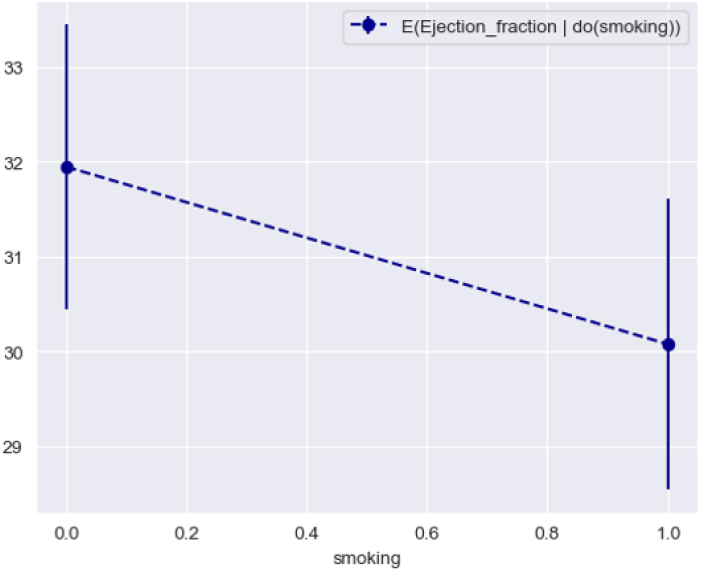
Visualization of Estimated ATEs of smoking on ejection fraction in heart failure patients.

**Fig. 16.**
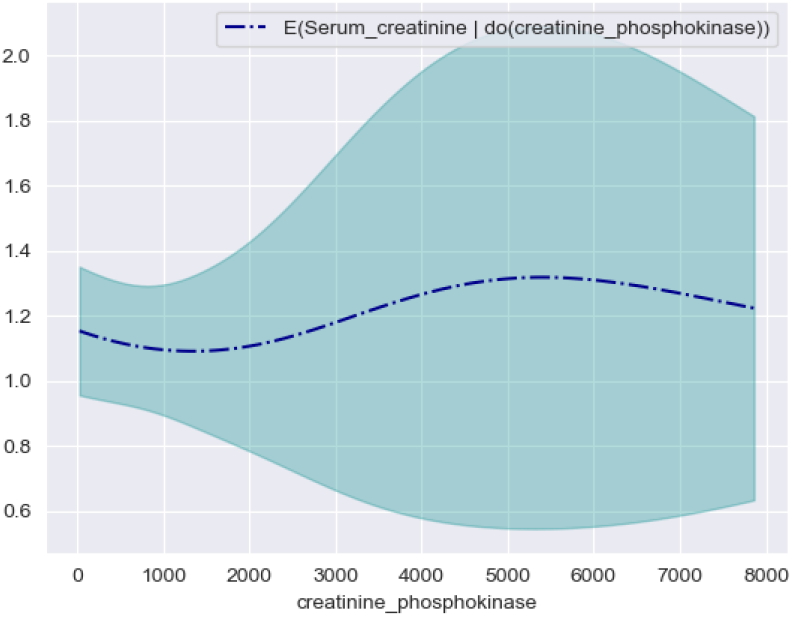
Visualization of Estimated ATEs of creatine phosphokinase on serum creatine in heart failure patients.

Meanwhile, in creatine phosphokinase(CPK) intervention results, three distinct ATE intervals were found to exist: [0mcg/L,1000mcg/L], [1000 mcg/L, 5000 mcg/L], and [5000 mcg/L, 8000 mcg/L]. In interval: [1000 mcg/L, 5000 mcg/L], intervention in CPK was found to increase average serum creatine levels, whereas in intervals: [0 mcg/L,1000 mcg/L] and [5000 mcg/L, 8000 mcg/L], weak negative(-) intervention effects were found to be present. However, as CPK was increased through intervention, standard deviations of ATE exhibited drastic increase, which led to the fact that changes found in average intervention effects in current research are not sufficient to represent precise causal relationships between CPK and serum creatine. Thus, it was found that indirect results from intervening in CPK levels should be analyzed under advanced conditions where additional confounders are being considered under precise domain knowledge.

## VI. Conclusion

In this paper, based on the concepts of causal DAG extraction using Graph AutoEncoders and NCM learning with Structured neural network (StrNN), deep causal inference and intervention analysis regarding risk factors of death in heart failure patients and mortality were sequentially implemented. Results show that under valid causal relationships extracted from GAE, StrNN based causal models can return significantly better mortality predictions than conventional ML based models (such as Random Forest or SVM radial models) in terms of prediction accuracy, F1-scores and Matthews Correlation Coefficients, even though less data is used for fitting. Especially, StrNN with batch size=50 outperformed results from ML based feature selection + fitting in both test accuracy and MCC by approximately 17.3%p and 5.96%p.

Under GAE and StrNN, three risk factors were extracted as direct causal factors of death in heart failure patients: Serum Creatinine(mg/dL), Platelets(kiloplatelets/mL) and Ejection fraction(%). Validity and effectiveness of using StrNN models being checked, intervention analysis for each factor was computed under do-calculus rules and Pearl, J’s backdoor criterion. From estimated ATE results for each causal factor, three significant findings were deduced. First, in serum creatinine ATEs, it was found that interventions to decrease serum creatinine levels within interval of [0mg/dL, 4mg/dL] can lead to drastic alleviations of death probabilities in heart failure patients. Meanwhile, it was also found that interventions to decrease serum creatinine levels within the interval of [4mg/dL, 8mg/dL] would not lead to alleviations of death probabilities, since ATE becomes invariant to serum creatinine level changes in interval [4mg/dL, 8mg/dL]. Second, in platelet ATEs, though positive(+) causal relationships were found in several intervals, it was found that decrease of platelets within intervals: [0 kiloplatelets/mL, 80000 kiloplatelets/mL], [100000 kiloplatelets/mL, 320000 kiloplatelets/mL] and [340000 kiloplatelets/mL, 800000 kiloplatelets/mL] does not lead to significant changes in ATE regarding death event probabilities in heart failure patients. Lastly, in ejection fraction ATEs, it was found that interventions to increase ejection fractions within interval [25%,45%] can lead to significant decrease in death event probabilities in heart failure patients. Meanwhile, in intervals, [0%, 25%] and [45%, 80%], ATE was found invariant to interventions or changes in ejection fraction percentage.

Through results from this research, it is expected that better clinical approaches can be implemented to prevent heart failure patient mortalities who had left ventricular systolic dysfunction or who were classified as class III or IV in NYHA criterion due to heart failure. Furthermore, as deep causal modelling based on GAE and StrNN was found valid and effective in analyzing observation data regarding CVD patients, vast usage or incorporation of fitted StrNN model results in predicting death events among heart failure patients is expected to highly increase survival rates within patients who are at risk of death due to CVD related factors.

Though significant findings were deduced under this research, there were also certain limitations in terms of sample size, ethnicity and hyper parameter settings. As the implemented data: “UCI repository Heart Failure Clinical Records data” is a small-sample dataset, causal effects from other significant risk factors might have been overlooked. Meanwhile, as the patient’s ethnicity was substantially limited, generalization of results from research to other ethnic groups might have slight discrepancies, when compared to ground-truth knowledge. Lastly, due to the existence of numerous hyperparameters in GAE settings, there were certain limitations in considering or probing for optimal hyperparameter sets for causal DAG extraction. In further analysis or follow-up studies, it would be ideal to solve these limitations to construct better, high performing NCMs with stronger validity and universality.

